# *In vivo* modulation of locus coeruleus activity by light

**DOI:** 10.64898/2026.04.28.721313

**Authors:** Fermin Balda Aizpurua, Roya Sharifpour, Islay Campbell, Elise Beckers, Sara Lelot, Alexandre Berger, Nasrin Mortazavi, Ekaterina Koshmanova, Fabienne Collette, Laurent Lamalle, Heidi IL Jacobs, Mikhail Zubkov, Puneet Talwar, Gilles Vandewalle

**Affiliations:** GIGA-Institute, CRC-Human Imaging, University of Liège, Liège, Belgium; Faculty of Health, Medicine and Life Sciences, School for Mental Health and Neuroscience, Alzheimer Centre Limburg, Maastricht University; Maastricht, The Netherlands; Athinoula A. Martinos Center for Biomedical Imaging, Department of Radiology, Massachusetts General Hospital and Harvard Medical School, Boston, MA, USA; Department of Translational Neuroimaging, Hôpital Erasme, Hôpital Universitaire de Bruxelles, Université libre de Bruxelles, Brussels, Belgium

**Keywords:** non-image-forming (NIF) light, melanopsin, hypothalamus, norepinephrine, Ultra high-field MRI

## Abstract

Dysfunction of the Locus coeruleus (LC), a small brainstem nucleus which consists of the brain’s primary source of norepinephrine, is linked to various neurological and psychiatric disorders. Light exposure influences many non-image-forming (NIF) biological functions including those modulated by LC activity. Yet direct evidence of light-driven modulation of the LC has remained elusive due to the structure’s small size and deep location. Using ultra-high-field (7 Tesla) functional MRI during an emotional task under varying illuminances, we demonstrate that light modulates LC activity in an emotion-dependent manner. We show that increased illuminance dampens LC responses to negative emotional stimuli while enhancing responses to neutral stimuli. Furthermore, we provide tentative evidence that within an emotional context, light may affect LC activity through the basomedial nucleus of the amygdala as well as through a region of the hypothalamus encompassing the lateral hypothalamus. These findings open avenues for the development of targeted, non-pharmacological treatments leveraging light to modulate norepinephrine tone and/or LC activity in several neuropsychiatric disorders.

**Significance Statement:** Dysfunction of the Locus Coeruleus (LC), a small nucleus of brain and the main source of norepinephrine, is linked to various brain disorders. Here we provide in vivo evidence that exposure to light can alter the activity of the LC with higher illuminance leading to dampened LC responses to negative emotional stimuli. These findings identify the LC as a key target for the therapeutic effects of light on emotion, offering a biological framework for using light to treat neuropsychiatric disorders.

## Introduction

The locus coeruleus (LC) is a small nucleus of the brainstem - it measures approximately 12 to 17 mm in length, roughly comparable to the diameter of a penny - that holds great promise as an intervention target for many neurological conditions. The LC is the main source of norepinephrine in the brain and has widespread monosynaptic projections to all cerebral territories, including regions essential to sleep, cognition and mood (1). Interest in the LC is notably rising because it is among the first brain sites showing neuropathological signs of neurodegenerative diseases, such as Alzheimer’s disease (AD) and Parkinson’s disease (PD), decades before the typical onset of the symptoms. Furthermore, changes in LC MRI integrity have been related to early Alzheimer’s disease-related cognitive symptoms (2). Dysfunction of the LC may also contribute to the emergence of insomnia disorder, the second most common psychiatric disorder worldwide (3, 4). In addition, altering LC function, through vagus nerve stimulation (VNS) or pharmacological interventions, may improve memory function in healthy older individuals, in Alzheimer’s disease and epilepsy (2, 3, 5–7), while reducing the severity of seizures in patients with refractory epilepsy (8). Furthermore, norepinephrine neuromodulation is the target of many anti-depressant drugs (9). Aside from these interventions, research on animal models and in humans has provided indirect evidence that the LC activity is acutely affected by light exposure (10–13). Light could therefore constitute a simple intervention to affect LC-dependent functions, resulting in positive effects on cognitive, mood and sleep disorders (14–16).

Exposure to light is required for the fine-tuning of numerous non-image-forming (NIF) biological processes, including aspects that depend in part on LC activity, such as the regulation of cognition, anxiety, mood, alertness and sleep (17–19). At the retinal level, the NIF effects of light heavily depend on intrinsically photosensitive retinal ganglion cells (ipRGCs) which express the blue-wavelength-sensitive photopigment melanopsin (10). Depending on light illuminance, ipRGCs combine the inputs of rods and/or cones with their intrinsic melanopsin-driven responses to funnel light signal to the brain with a larger efficiency around the blue portion of visible light (20, 21). IpRGCs target multiple, mostly subcortical, brain structures, especially within the hypothalamus (10, 19). Critically, magnetic resonance imaging (MRI) was previously used to report light-induced modulation of a dorsal part of the pons, but the neuroimaging procedure did not allow ascertaining whether it was within the LC (13). Likewise, the transient pupil dilation associated with a salient emotional stimulus, a response that is in part mediated by the LC (22), was affected by light illuminance (23). Beyond this indirect evidence, whether ambient light affects LC activity has not yet been established in animals or in humans, in part because of the difficulty of imaging such a small and deep structure.

Here, we tested whether light illuminance affects the responses of the LC to emotional and neutral auditory stimuli in healthy young adults of both sexes. We leveraged ultra-high-field 7 Tesla MRI to precisely delineate the LC – using an LC-dedicated MRI sequence (24) – and extract its activity while exposed to varying experimental light conditions. We further explored linear relationships between the modulation of activity by light of the LC and of subparts of the amygdala and hypothalamus. These regions were selected as they are key nodes in the NIF pathway that exhibit high sensitivity to light (12, 25) in humans and project directly or indirectly to the LC in animal models (26–28). We anticipated that, compared to neutral items, the responses of the LC to emotional stimulation would be particularly modulated by light illuminance – but did not have a priori hypothesis regarding the direction of this modulation.

## Results

Thirty-five healthy young participants (22 women; 24.5 ± 3.3y; **Supplementary Table S1**) completed an fMRI protocol conducted at University of Liège (Belgium) in the morning (3 to 3.5hrs after habitual wake-up time) including an emotional auditory task that was previously successfully used to demonstrate that light quality affects emotional brain responses (29, 30) (together with 2 other cognitive tasks not included in the present analysis; **Figure 1A-C**). During the task, participants stayed in darkness or were exposed to different light consisting of a blue-enriched polychromatic white light of three different illuminance levels (37, 92, 190 melanopic Equivalent Daylight Illuminance – melanopic EDI - lux) and a low illuminance monochromatic orange light (0.16 melanopic EDI lux) (**Supplementary Table S2**).

**Figure 1.**
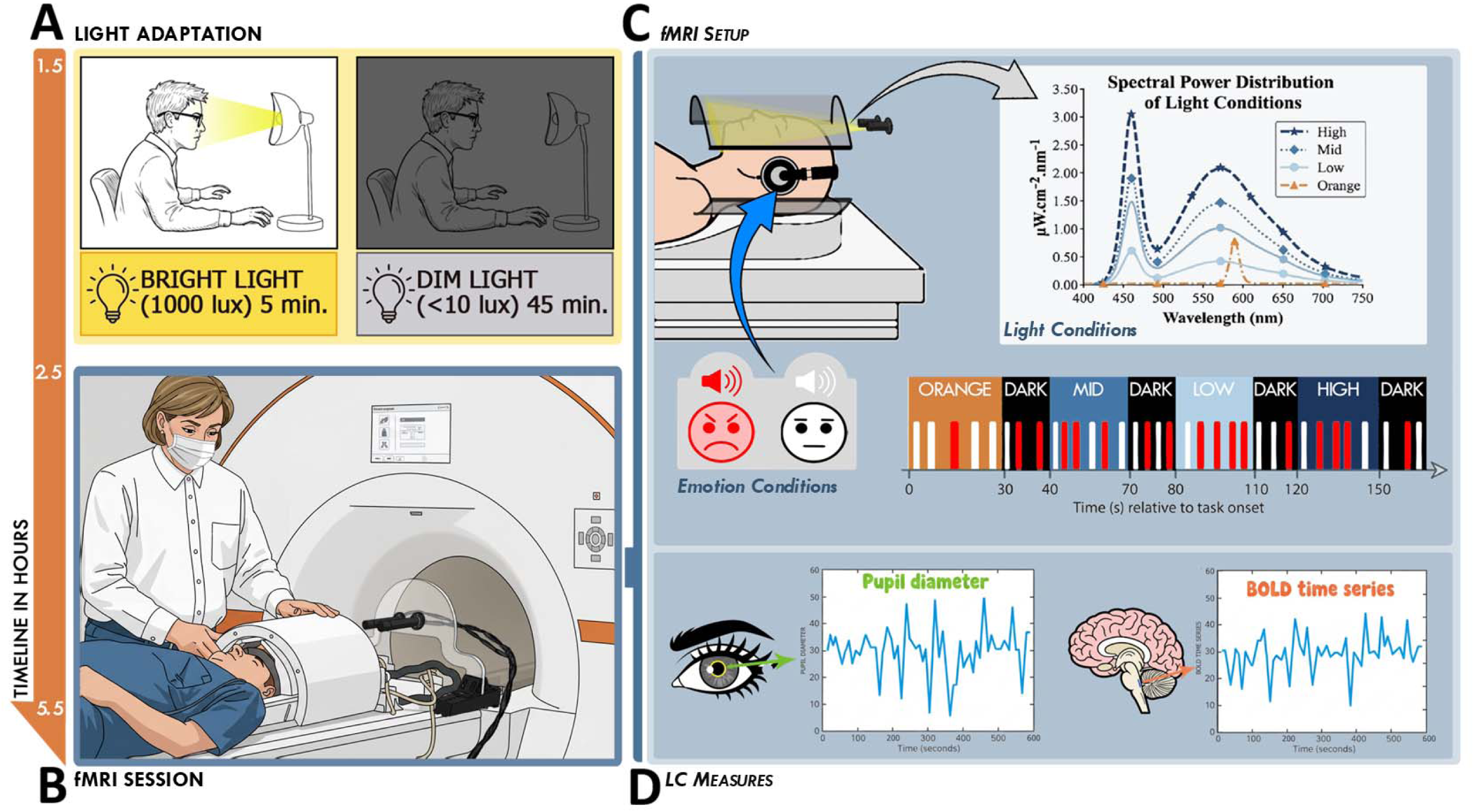
Experimental protocol. **(A) Light Adaptation.** Following 7 days of actigraphy-verified regular sleep-wake schedule (± 1h), participants arrived at the laboratory 1.5h after their habitual wake-up time. They underwent a light adaptation protocol consisting of 5 minutes of bright light adaptation (1000 lux) followed by 45 minutes of dim light (<10 lux) adaptation before entering the fMRI scanner. **(B) fMRI Session.** Within the session, 35 participants (24.49 ± 3.32 y; 22 women) performed an executive task, an emotional task, and an attentional task. Only the emotional task is analyzed in this manuscript. **(C) fMRI Set Up.** Participants were positioned with their heads inside a 7T MRI coil. Light was delivered via an optic fiber to illuminate the inside of the coil, exposing participants to four light conditions defined by their Spectral Power Distribution (SPD): Monochromatic orange (0.16 mel EDI lux) and three polychromatic blue-enriched levels (6500K; Low: 37 mel EDI; Mid: 92 mel EDI; High: 190 mel EDI; see **Suppl. Table S2**). Blue-enriched illuminances were selected to maintain photon flux consistency with prior studies (29, 30). While under these light conditions, participants performed an emotional task involving gender classification of meaningless vocalizations (32). Unknown to participants, 50% of vocalizations were neutral (white) and 50% were pronounced with angry/emotional prosody (red) by professional actors. For current ROI analyses, conditions were indexed solely by melanopic EDI lux. Panels A-C copied from under a CC BY 4.0 license (12). **(D) LC Measures.** Continuous pupillometric data were recorded using an MR-compatible infrared eye-tracker. Pupil diameter is a well-established non-invasive proxy for central norepinephrine (NE) activity. Whole-brain BOLD signal was also recorded during the task, allowing activity extraction of the LC.

Participants achieved 92 ± 6.99% (mean ±SD) button response, while the accuracy to the lure sex classification task (31) was 79.21% ± 9.10%, which is slightly lower than previous studies using the same task (32). In line with previous literature (30, 32), the reaction times were faster for the neutral as compared with the emotional stimuli (main effect, stimulus type; *F*(1, 23) = 15.661, ***p* < .001**, η^2^ = .405) (**Figure 2A**), confirming that the emotional content of the stimuli was successful in triggering a behavioural response. Importantly, reaction times exhibited a marginal main effect of illuminance (main effect, illuminance; *F*(4, 92) = 2.408, *p* = .054, η^2^ = .094) suggesting a trend toward accelerated processing speeds as illuminance increased (**Figure 2B**). In contrast, there was no interaction between illuminance and stimulus type (*F*(4, 92) = 0.389, *p* = .816) and none of the other covariates were significant (**Supplementary Table S3**).

**Figure 2.**
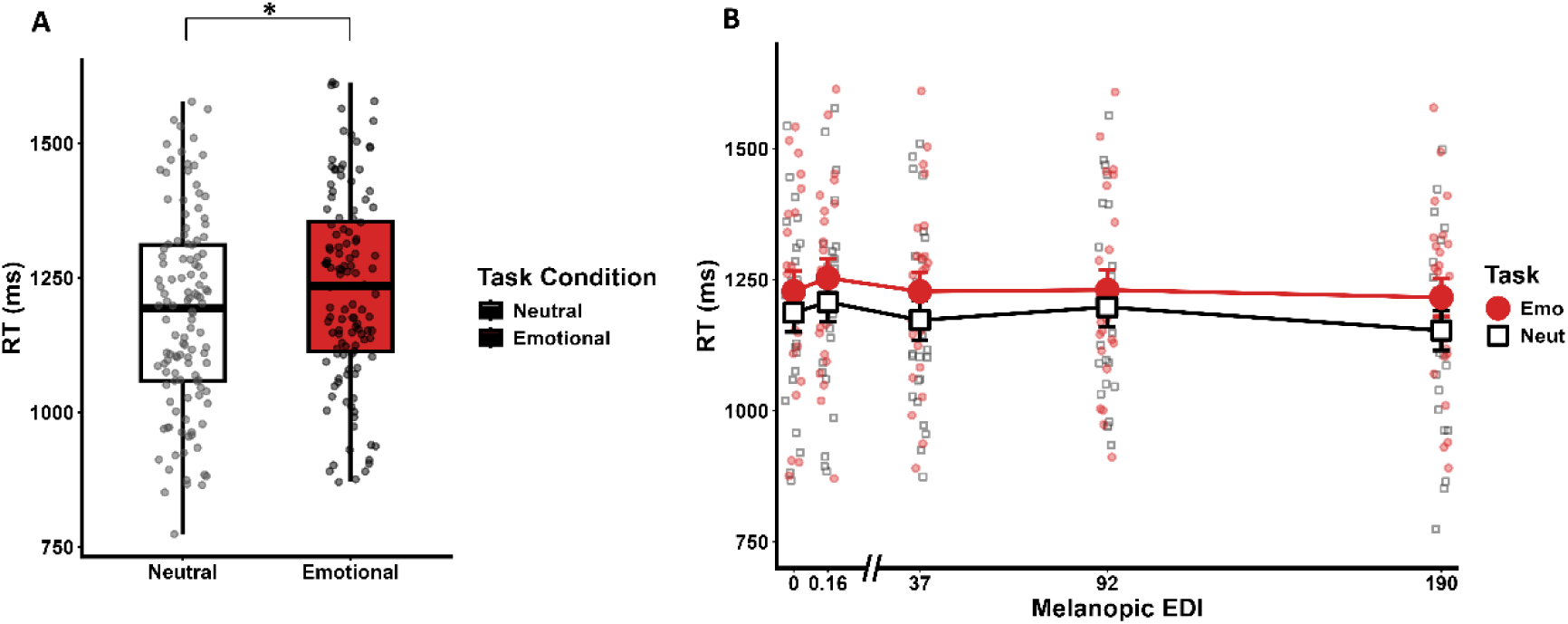
Impact of Light on Task Performance. **(A) Impact of emotional prosody on reaction times.** Reaction times (ms) were faster for neutral vocalizations compared to emotional (angry) vocalizations (p <. .001), confirming that the emotional content effectively modulated behavioral responses. **(B) Light-induced modulation of reaction times.** A nominally significant main effect of illuminance was observed (p = .055), with reaction times becoming progressively faster as light illuminance increased across conditions. No significant interaction was found between illuminance and stimulus type for reaction time.

Expert delineation of the LC on MR-images providing contrast to identify the LC anatomy (33) (**Figure 3A**), enabled the extraction of the Blood Oxygen Level Dependent (BOLD)-responses in the LC to the neutral and emotional stimuli. The first statistical analysis revealed that the LC was differentially recruited by stimulus type (*F*(1, 34) = 4.578, ***p* = .0396**, η^2^ = .118) (**Table 1**; **Figure 3C**) with no contribution from age, sex, BMI and season, included as covariates. Critically, our main statistical model, focused on the overall impact of light on LC responses (and including the same covariates), also yielded a significant difference between task conditions revealing that light illuminance differently affected the responses of the LC to the neutral and emotional stimuli (*F*(1,34) = 13.947, *p* < .001, η^2^ = .291) (**Table 1**; **Figure 3D**). Increased illuminance was associated with progressively lower and higher responses of the LC activity to the emotional and neutral stimuli, respectively. The whole brain group analysis (p_uncorrected_ <.05) yielded local positive peaks bilaterally in the LC supporting that our finding does not arise from partial volume effects or ‘leaking’ activation from nearby structures or the adjacent 4th ventricle (**Figure 3B**).

**Figure 3.**
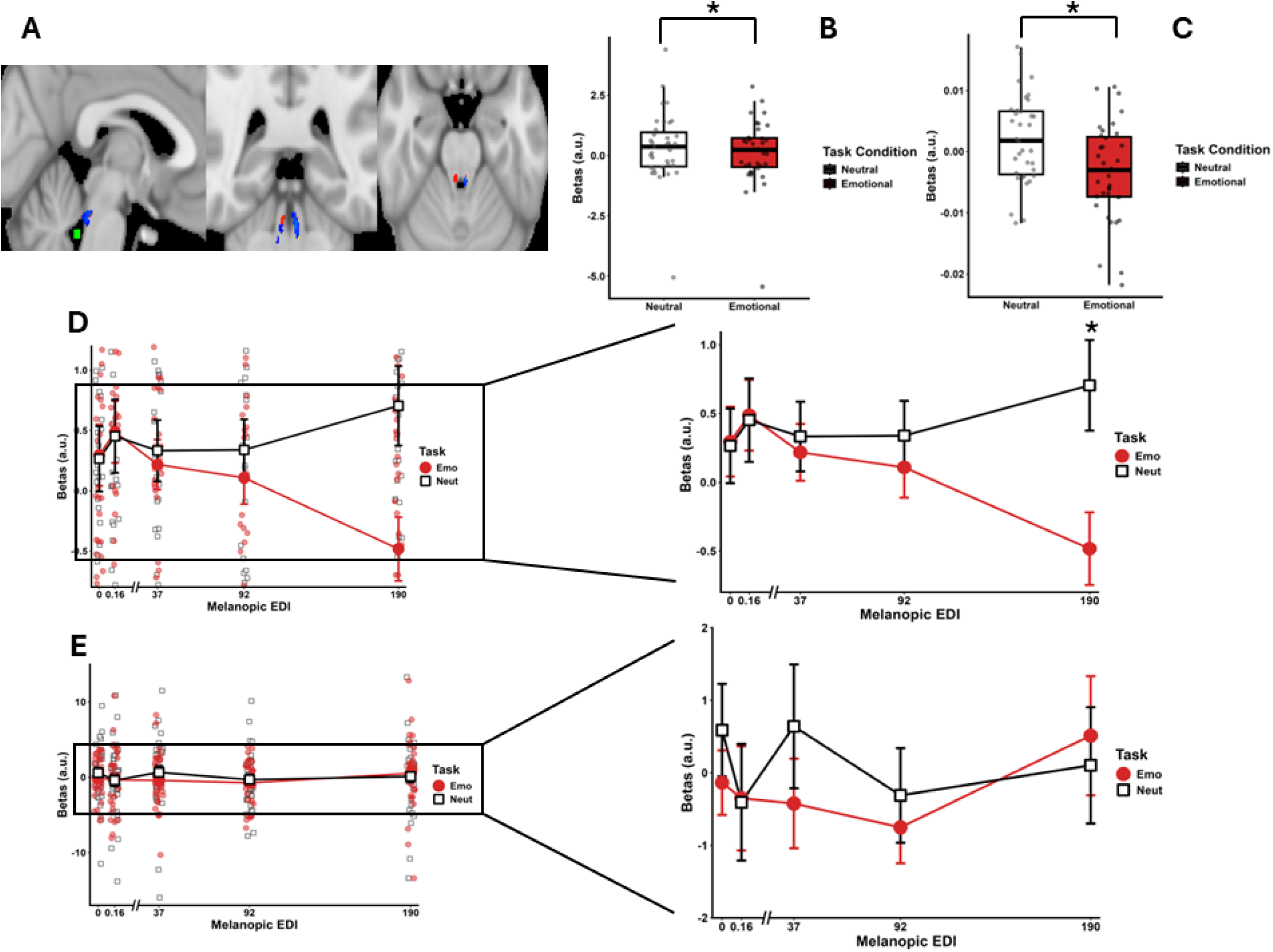
Modulation of Locus Coeruleus activity by illuminance during emotional and neutral processing. **(A) Functional cluster localization.** Axial, sagittal, and coronal views showing the group-average LC mask (blue) and the mask of the 4th ventricle (green), used to verify the spatial specificity of the findings. Voxel-wise group results (red) identify clusters within the LC where the contrast between emotional and neutral stimuli was significantly modulated by light illuminance with a local maxima within the LC (p_uncorrected_< .05). Group results were computed for visualization only, following normalization to MNI152 space (activity estimated used in the other panels were extracted within subject space) to ensure that our findings do not arise from a nearby ‘leaking’ activation/deactivation. **(B) Endogenous LC recruitment.** Estimates (beta; arbitrary units – a.u.) of LC activation during emotional (Emo) and neutral (Neut) task blocks irrespective of the light condition (Refer to **Table 1** for full statistics). **(C) Light-modulated LC recruitment.** Estimates of the parametric modulation of LC activity by melanopic EDI illuminance (Refer to **Table 1** for full statistics). Boxplots illustrate LC recruitment as a function of illuminance (0.16, 37, 92, and 190 lux). A significant Light-by-Task interaction was detected (p < 0.001), adjusted for age, gender, BMI, and seasonal variation. **(D) Simple effects of task across light intensities in LC.** Comparison of task-related LC activation at each illuminance. (Left) Full-scale plot including individual data points to illustrate inter-individual variability. (Right) Zoom-in of the same plot to highlight mean differences. Tukey-adjusted post-hoc comparisons revealed that neutral stimuli elicited significantly greater LC recruitment compared to emotional stimuli at both 92 lux (p = 0.037) and 190 lux (p < 0.001) (refer to Table 2 for full statistics). **(E) Simple effects of task across light intensities in 4^th^ ventricle.** Comparison of task-related BOLD Signal within the 4^th^ ventricle at each illuminance. (Left) Full-scale plot including individual data points to illustrate inter-individual variability. (Right) Zoom-in of the same plot. No significant main effect of light or task or interaction (refer to Table S3 for full statistics).

**Table 1.** Summary of GLMM results comparing baseline and light-modulated models of LC activity. F-statistics are reported for fixed effects with degrees of freedom (df) in parentheses, estimated using the Satterthwaite approximation. *P < .05, **P < .01, ***P < .001.

| Emotional Task LC Analysis |  |  |  |
| --- | --- | --- | --- |
| Effect | *F* (*df*) | *p* | $\eta^2$ |
| Fixed Effects (Baseline Model) |  |  |  |
| Task | 4.58 (1,34) | **0.040\*** | 0.119 |
| Age | 3.08 (1,30) | 0.090 | 0.093 |
| Gender | 0.33 (1,30) | 0.572 | 0.011 |
| BMI | 0.43 (1,30) | 0.515 | 0.014 |
| Season | 0.46 (1,30) | 0.502 | 0.015 |
| Fixed Effects (Lightmod Model) |  |  |  |
| Task | 4.58 (1,34) | **<0.001\*\*\*** | 0.291 |
| Age | 0.04 (1,30) | 0.831 | 0.002 |
| Gender | 0.31 (1,30) | 0.581 | 0.010 |
| BMI | 1.48 (1,30) | 0.233 | 0.047 |
| Season | 0.44 (1,30) | 0.509 | 0.015 |

**Table 2.** GLMM ^r^esults for the factorial analysis of light and emotional task effects on LC activity. F-statistics are reported for fixed effects with degrees of freedom (df) in parentheses, estimated using the Satterthwaite approximation. Pairwise comparisons (contrasts) for the Light x Task interaction represent the difference between Emotional and Neutral conditions (Emo–Neut) across light levels. t-values represent the strength and direction of these contrasts. *P < .05, **P < .01, ***P < .001.

| Emotional Task LC Analysis Factorial |  |  |  |
| --- | --- | --- | --- |
| Effect | *F* (df) | *p* | $\eta^2$ |
| **Fixed Effects** |  |  |  |
| Light | 0.48 (4, 133) | 0.753 | 0.014 |
| Age | 3.30 (1, 30) | 0.079 | 0.099 |
| Gender | 0.36 (1, 30) | 0.554 | 0.012 |
| BMI | 0.37 (1, 30) | 0.547 | 0.012 |
| Season | 0.57 (1, 30) | 0.457 | 0.019 |
| Task | 5.58 (1, 34) | **0.024 \*** | 0.141 |
| Light x Task | 4.07 (4, 133) | **0.004 \*\*** | 0.109 |
| **Posthoc Comparisons** |  |  |  |
| Contrast | *t* | *p* |  |
| (Emo-Neut) Dark | 0.125 | 0.901 |  |
| (Emo-Neut) Orange | 0.206 | 0.837 |  |
| (Emo-Neut) B1 | -0.443 | 0.658 |  |
| (Emo-Neut) B2 | -1.342 | 0.182 |  |
| (Emo-Neut) B3 | -4.512 | **<.001\*\*\*** |  |
| Emo (0 vs 190 lux) | 2.431 | **0.016\*** |  |
| Emo (0.16 vs 190 lux) | 2.916 | **0.004\*\*** |  |

Only significant pairwise comparisons are reported in the main text. For all comparisons, refer to Table S4

When considering in more detail the activity of the LC under each illuminance, our statistical analysis confirmed an interaction between illuminance and task condition (*F*(4, 136) = 3.118, ***p* =.0172**, η^2^ = .084) (**Table 2**; **Figure 3E**), still including age, sex, BMI and season as covariates. Post hoc comparisons showed that LC responses to emotional trials (190 > 0.16 & 0 melanopic EDI lux, p_uncorrected_<0.016), with a significant difference between neutral and emotional stimuli at 190 melanopic EDI lux (p_corrected_ < 0.001) (**Table 1 and Supplementary Table S4**).

Given the small size of the LC, we further conducted control analyses to support our findings. As a negative control we extracted BOLD signal from the 4^th^ ventricle, near the LC, where one would not expect light to have any impact (**Figure 2A** for voxel location). The statistical analysis confirmed the absence of effect of the stimulus type and illuminance (**Figure 2F; Supplementary Table S5**). As previously (23), we further confirm that transient pupil dilation associated with the auditory stimuli was stronger under higher illuminance (*F*(1, 90.639) = 9.235**, *p* < .001**, η^2^ = 0.290), further supporting that light affects LC activity (in participants with usable data; N = 24) (**Supplementary Table S3, Supplementary Figure 2**). Finally, we examined correlations between task performance or transient pupil dilation and LC responses and found no significant associations (**Supplementary Table S6**).

Building upon previous reports that light exposure modulates emotional responses in the hypothalamus and amygdala in a subpart of the present dataset (N=26 and N=29) (12, 25), we explored whether the overall impact of illuminance on these regions was correlated with its impact on the LC. We reported an anterior-posterior gradient for the impact of illuminance within the hypothalamus (12) and therefore, we now focused on the anterior inferior subpart, encompassing the suprachiasmatic nucleus (SCN) and supraoptic nucleus (SON), and the posterior lateral part encompassing the lateral hypothalamus (LH) and tuberomammillary nucleus (TMN). Separate analyses for emotional and neural items (and including covariates) resulted in no significant association between LC activity and the anterior hypothalamus, while a significant correlation was detected for emotional stimuli with the posterior/lateral hypothalamus (t=1.798, **p=0.037**; **Figure 4; Supplementary Table S7**). We then considered the Basomedial nuclei and Medial and Cortical nuclei subparts of the amygdala as they showed the most prominent impact of light illuminance (25). Separate analyses for emotional and neural items led to a nominally significant association with the Basomedial nuclei subpart for emotional stimuli (t = 2.08, **p = 0.046**) and no association with the Medial and Cortical nuclei subparts (**Figure 4; Supplementary Table S7**). These explorations suggest that the local impact of illuminance on the activity of the LC may be related to local activity within the amygdala and within the hypothalamus for emotional but not for neutral items.

**Figure 4.**
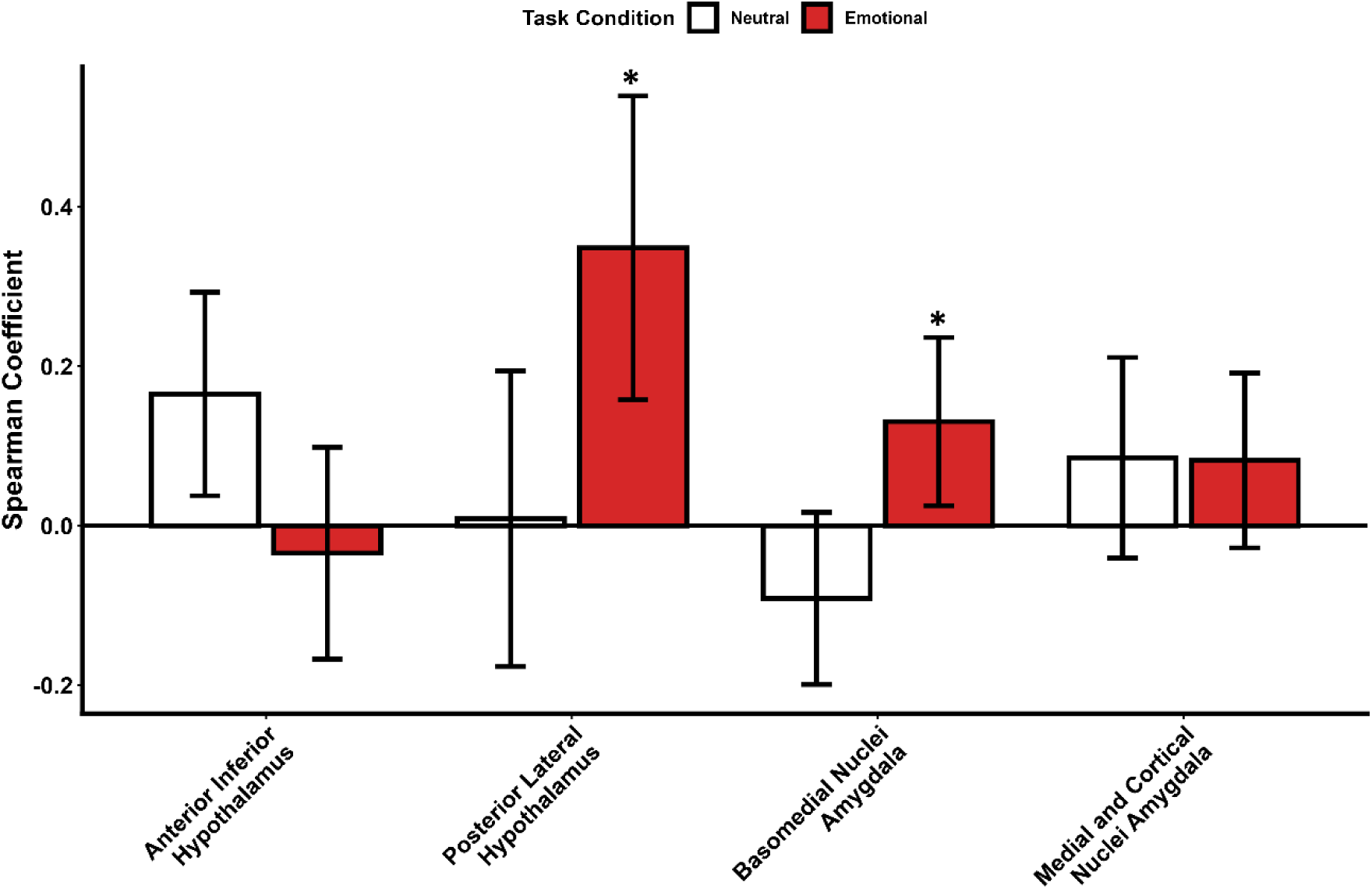
Comparative task-related correlation between the Locus Coeruleus (LC) and subcortical regions of interest. Bar plots illustrate the correlation strength (expressed as Spearman coefficients) between stimulus-evoked responses of the LC and specific subparts of the hypothalamus and amygdala: Anterior inferior hypothalamus, Posterior lateral hypothalamus, Basomedial nuclei of the amygdala, and Medial and Cortical nuclei of the amygdala. White bars represent the LC correlation during neutral stimulus processing, and red bars represent the correlation during emotional stimulus processing. Error bars denote the standard error (SE) of the linear regression slope. These coefficients represent the relationship between regional activities and do not substitute for the GLMM outputs reported in the main text (Full statistical output in Table S7).

## Discussion

The LC is attracting interest as a potential intervention target for neurological and psychiatric disorders (34). While pharmacology and VNS stand as promising means to stimulate the LC (35), here, we report that modulating activity of the LC while executing a cognitive task can be achieved by modifying the quality of light exposure. We show that the activity of the LC in response to auditory emotional stimuli progressively decreases with increasing illuminance, while the pattern is reversed for neutral stimuli. We further provide evidence that the impact of light on the LC may increase the coupling between the LC and the posterior lateral part of the hypothalamus, notably the LH and TMN, as well as the basomedial nuclei of the amygdala. These findings provide more insight into the neural mechanisms underlying light-induced changes in brain function and may have implications for the design of interventions for brain disorders using light or mimicking light effects.

The relationship between behavior and the functioning of the LC is considered to follow an inverted-U shape dynamic (36, 37). Accordingly, optimal arousal and cognitive performance is achieved when tonic activity of the LC is at intermediate level. This level would allow relatively pronounced transient responses of the LC to incoming salient stimulus, whereas low or excessively high tonic activity is associated with drowsiness or distractibility, respectively (36, 37). Our results suggest that light affects the activity of the LC in a stimulus-type-dependent manner. Specifically, as light illuminance increases, the LC’s response to the more salient negative stimuli is progressively dampened. This could be the result of a reduction in the phasic response of the LC or a progressive increase in the tonic activity of the LC, reducing its phasic expression. In contrast, the response to the less salient neutral stimulation would be progressively increased with increasing illuminance, as reported for other non-emotional items in different cognitive contexts [e.g. (13)].

We did not include positive emotional stimulation in the protocol – positive emotions are more difficult to trigger consistently across participants (38) - so we do not know whether positive stimuli, which are also salient and presumable affected by alertness and the level of tonic activity of the LC, trigger an enhanced response of the LC with increasing illuminance. Given the accepted positive impact of light therapy (39), we interpret our findings as a reflection of a positive impact of light on emotional brain function, which could reduce responses to negative stimulations. Importantly, a control analysis of the adjacent 4th ventricle elicited no significant changes in BOLD signal, confirming that our results do not stem from non-specific physiological noise. We further emphasize that the impact of light we report is of limited effect size (η² = .291 corresponding to medium effect size). While our protocol focused on the immediate effects of transient light exposure, we hypothesize that sustained and repeated exposure of such modulatory effects over weeks contributes to the sustained clinical improvements in affective state associated with long-term light therapy.

Animal data did not report direct projections from ipRGCs to the LC. However, there are indirect projections from the ipRGC targets to the LC. The SCN, site of the main circadian clock, receives dense ipRGC inputs and indirectly projects to the LC through the dorsomedial hypothalamus (DMH) (11, 40). The orexinergic LH also received ipRGC inputs and projects to the LC to regulate arousal and the maintenance of wakefulness (41). Here, we find a statistical trend that the activity modulation of the LC by light illuminance is correlated to the activity of the posterior subpart of the hypothalamus. We therefore speculate that within the context of our protocol, it may be the projection from the LH to the LC rather than the projection from the SCN to the LC that mediates the impact of light on LC activity. We also find a nominally significant correlation between LC and the basomedial nuclei, but not with the medial and cortical amygdala nuclei even though IpRGCs directly project to these latter nuclei. It is possible that light signal is transmitted to the LC via an indirect projection to the basomedial amygdala, but this requires further investigation, using for instance advanced connectivity analyses, such as dynamic causal modeling (DCM) and high resolution tractography (10, 23)

We further note that while a seasonal variation in the activity of the amygdala and in the impact of light illuminance on its activity was observed (25), we did not find seasonal changes in LC activity in the present study. We assume therefore that in contrast to other structures of the brain (42) LC activity is not showing prominent seasonal variations, at least within our experiment set up. In addition, even though reaction times to the auditory stimuli were affected by light illuminance, we did not find significant correlation between the activity of the LC (or of the amygdala like previously reported (25)) and reaction time. This is potentially a reflection of the fact that behavior is primarily mediated by the prefrontal cortex, a region that receives dense noradrenergic projections from the LC, and that a network of cortical and subcortical areas may be jointly affected by light to set behavior.

In addition, while we conducted our investigation in the morning, which corresponds to the commonly recommended time-of-day for light therapy, we did not vary the time-of-day of light administration when it is known to influence the impact of light (43, 44). This could also be addressed in future investigation, together with other factors including prior sleep duration and age (45, 46). Future research could include longitudinal assessments prior and after a light intervention in patients suffering from Seasonal Affective Disorder (SAD), to confirm that light can positively influence affect through the LC. The acute effects we report may be relatively independent from the longer-term effect of light therapy and represent a direct modulation of brain function (47). In addition, we only considered illuminance variation as our variable of interest and ignored the colour difference between the lowest illuminance (i.e. orange) and the others (i.e. blue enriched). Therefore, we cannot determine whether colour contributes to our findings. In addition, while the involvements of the melanopsin photopigment and of ipRGCs are very likely given the light level we used, we cannot confirm them. IpRGC integrate their intrinsic melanopsin photoreception with those of rods and cones that were also stimulated by the light (1). Rods and cones further project to many other types of RGC that also send light signaling to the brain, mostly however toward vision-related areas – in contrast to ipRGCs. Future studies should work with other light spectra or with metameric light exposures and silent substitution paradigms (48) to disentangle whether LC modulation arises primarily from melanopsin-driven input or integrated photoreceptor signaling.

Light therapy is already established as an effective intervention for mood disorders such as SAD and non-seasonal depression (49, 50). Our findings provide novel insights into the neurobiological mechanisms through which light exposure may influence emotional processing, particularly by modulating activity within the LC, a key node in the brain’s arousal and affective regulation systems. Beyond depression, dysregulation of the LC–norepinephrine system is implicated in several neuropsychiatric disorders — including post-traumatic stress disorder (PTSD), and generalized anxiety disorder (GAD) — supporting the translational potential of light-induced modulation of LC activity (51, 52). In addition, since several classes of Selective Norepinephrine Reuptake Inhibitors (SNRIs) drugs specifically target the noradrenergic system to alleviate symptoms of depression and anxiety (53), our results provide a neurobiological rationale for the efficacy of light therapy. By modulating LC activity, light may act through similar pathways as these pharmacological interventions, offering a low-risk adjunct or alternative for mood regulation that typically presents fewer and less severe side-effects than traditional SNRIs. Furthermore, since the LC is a critical hub for attentional filtering and cognitive flexibility (36), our findings suggest that targeted light exposure could serve as a non-pharmacological intervention to bolster cognitive function, particularly maybe in demented populations (54) LC integrity is compromised and sleep-wake cycles are frequently disrupted.

## Materials and Methods

### Study Overview

The data analyzed in this study were collected as part of a larger research initiative, with methodological details previously reported (23, 55, 56). The study protocol was approved by the Ethics Committee of the University of Liège Faculty of Medicine (ref. 2020/11). All participants provided written informed consent and received financial compensation. Data acquisition occurred at the University of Liège between December 2020 and September 2023.

### Participants

35 young adults (22 women; 24.5 ±3.3y) were included in the analyses. Exclusion criteria were assessed through questionnaires and a semi-structured interview: history of psychiatric and neurological disorders, sleep disorders, use of psychoactive drugs or addiction; history of ophthalmic disorders or auditory impairments; colour blindness; night shift work during the last year or recent trans-meridian travel during the last 2 months; excessive caffeine (>4 caffeine units/day) or alcohol consumption (>14 alcohol units/week); medication affecting the central nervous system; smoking; pregnancy or breast-feeding (women); and counter indication for MRI-scanning. All participants had to score <18 on the 21-item Beck Anxiety Inventory (up to mild anxiety (57)), and <14 on the Beck Depression Inventory-II (up to mild depression; (58)), <12 on the Epworth Sleepiness Scale (59), and <8 on the Pittsburgh Sleep Quality Index (60). Questionnaires further assessed chronotype with the Horne-Östberg questionnaire (61) and seasonality with the Seasonal Pattern Assessment Questionnaire (62). By excluding individuals with scores indicative of moderate-to-severe symptoms, we minimized potential confounding effects of affective dysregulation on the emotional task outcomes.

### Overall Protocol

Structural brain images were acquired 1 to 2 weeks before the experiment, during a visit which served as habituation to the experimental conditions. Participants then followed a loose sleep-wake schedule (±1h from habitual sleep/wake-up time) for 7 days, to maintain realistic entrained life conditions and avoid excessive sleep restriction (verified using wrist actigraphy -AX3, Axivity, UK- and sleep diaries). Participants arrived at the laboratory 1.5 to 2h after their habitual wake time. To standardize participants’ recent light history, they were exposed to 5min of bright white light (1000 lux) and were then maintained in dim light (<10 lux) for 45min. During the dim light exposure, participants were given instructions about the fMRI cognitive tasks and completed practice tasks on a luminance-controlled laptop (<10 lux) **(Figure 1A)**. The fMRI sessions consisted of an executive task (25min), an attentional task (15min), and an emotional task (20min), which is the only task included in the present paper **(Figure 1B)**. Participants always completed the executive task first as it was the most demanding task. The order of the following two tasks was counterbalanced across participants. While in the MR-scanner, participants were asked to keep their eyes open and try not to blink too much during the cognitive tasks. An eye-tracking system (EyeLink 1000Plus, SR Research, Ottawa, Canada) was monitored for proper eye-opening during all data acquisitions.

### Light Exposure

We delivered light from a light box (SugarCUBE, Ushio America, CA, USA) to the MRI coil using an 8m optic fiber (1 inch diameter, Setra Systems, MA, USA). This configuration allowed uniform illumination in the machine (considering technical difficulties and limitations) and prevented direct light exposure to the eyes. At the same time, a stand positioned the dual ends of the optic fiber with reproducible fixation and orientation. In addition, we used a filter wheel (Spectral Products, AB300, NM, USA) and optical fiber filters (monochromatic narrowband orange filter - 589mn; full width at half maximum: 10 nm - or a UV highpass filter - 433–1650 nm) to create the light conditions of the experiment.

Because of the MRI’s ferromagnetic properties, we couldn’t take measurements at the scanner. Instead, we moved the MRI coil, the optic fiber, the stand, and the rest of the instrumentation into a dark room. We then simulated experimental conditions by positioning a sensor 2 cm from the coil’s mirror (mounted at eye level). For each light condition, we measured illuminance four times, recorded spectrum twice, and computed the average. The inability to control eye location is a noteworthy limitation of our protocol; however, we do not expect this limitation to significantly affect the analysis and interpretation of our data.

The illuminance and technical characteristics of blue enriched white light produced similar photon flux to previous experiments in our lab (29, 30); approximately 10^12^ - 10^14^ photons/m^2^s). The orange light was introduced as a control visual stimulation for potential secondary whole-brain analyses. For the present region of interest analyses, we discarded colour differences between the light conditions and only considered illuminance as indexed by melanopic EDI lux, constituting a limitation of our study. For the emotional task, we employed five distinct light conditions: three levels of blue-enriched white light with melanopic EDI values of 37, 92, and 190 lux, respectively; one orange light condition (melanopic EDI 0.16 lux; 590 nm, full width at half maximum [FWHM] 10 nm); and a darkness light condition (<0.1 lux). The cognitive task consists of 5 light blocks for each light condition which lasted 30-40s (average was 35s), and they had 10s of darkness (< 0.1 lux) rest in between (**Figure 1C**).

### Emotional Task

The task consisted of a sex-related voice discrimination task with emotional and neutral prosody auditory vocalizations (**Figure 1C**). The task was validated by previous behavioral assessments and that we successfully used it in prior experiment to assess light impact on emotional brain responses (12, 25, 29–32, 63). Participants were not informed that 50% of the auditory stimuli were emotional (i.e., spoken with angry prosody), while the remaining 50% were neutral. Professional actors (50% women) pronounced 240 auditory stimuli consisting of three meaningless words (‘goster’, ‘niuvenci’, ‘figotleich’). In addition, we matched the stimuli for the duration (750ms) and mean acoustic energy to avoid loudness effects. During each 30 - 40 s light block, we pseudorandomly delivered four angry and four neutral prosody tokens, each token delivered every 3 - 5s. The darkness periods between 2 light exposures contained two angry and two neutral stimuli that accounted for 80 distinct voice stimuli. In total, we distributed 160 distinct voice stimuli (50% angry; 50% neutral) across the four light conditions.

### Data Acquisition

The MRI data were acquired in a 7T MAGNETOM Terra MR scanner (Siemens Healthineers, Erlangen, Germany) with a 32-channel receive and 1-channel transmit head coil (Nova Medical, Wilmington, MA, USA). Dielectric pads (Multiwave Imaging, Marseille, France) were placed between the subject’s head and receiver coil to homogenise the magnetic field of Radio Frequency (RF) pulses.

Multislice T2*-weighted fMRI images were obtained with a multi-band Gradient-Recalled Echo - Echo-Planar Imaging (GRE-EPI) sequence using paraxial slice orientation (TR = 2340ms, TE = 24ms, FA = 90°, in-plane FoV = 224 mm × 224 mm, matrix size = 160 × 160, 86 axial slices, no interslice gap, slice thickness = 1.4 mm, bandwith = 1840 Hz/Px, GRAPPA acceleration factor = 3, multiband factor = 2). To avoid saturation effects, the first three scans were discarded. To correct for physiological noise in the fMRI data participants’ pulse and respiration movements were recorded using a pulse oximeter and a breathing belt (Siemens Healthineers, Erlangen, Germany). Following the fMRI acquisition a 2D GRE field mapping sequence to assess B0 magnetic field inhomogeneity with the following parameters: TR = 5.2ms, TEs = 2.26ms and 3.28ms, FA = 15°, bandwidth = 737 Hz/Px, matrix size = 96 × 128, 96 axial slices, slice thickness = 2 mm, acquisition time = 1:38 min, no parallel/multislice imaging, was applied.

For anatomical imaging, a high-resolution T1-weighted image was acquired using a Magnetization-Prepared with 2 RApid Gradient Echoes (MP2RAGE) sequence: TR = 4300ms, TE = 1.98ms, FA = 5°/6°, TI = 940ms/2830ms, bandwidth = 240 Hz, matrix size = 256 × 256 x 224, GRAPPA acceleration factor = 3, voxel size = (0.75 x 0.75 × 0.75) mm3.

To further characterize the LC, a high-resolution 3D Magnetization Transfer (MT) weighted Turbo Flash sequence was performed (20 magnetization transfer pulses, TR = 400 ms, TE = 2.55 ms, FA = 350°, bandwidth = 300 Hz/Px, voxel size = (0.5 x 0.5 x 0.5) mm^3^, frequency offset = 2000 Hz). The protocol utilized Phase Partial Fourier and a centric k-space ordering. The MT-weighted acquisition (MT-ON) used 2 averages. This sequence was tilted to be perpendicular to the brainstem to optimize the visualization of the LC.

### Data Processing

We removed the background noise for MP2RAGE images, the background using an extension (https://github.com/benoitberanger/mp2rage) of Statistical Parametric Mapping 12 (SPM12; https://www.fil.ion.ucl.ac.uk/spm/software/spm12/) under Matlab R2019 (MathWorks, Natick, Massachusetts) (64). We then reoriented images using the ‘spm_auto_reorient’ function (https://github.com/CyclotronResearchCentre/spm_auto_reorient; Cyclotron Research Centre, 2015) and corrected for intensity non-uniformity using the bias correction method implemented in the SPM12 ‘unified segmentation’ tool (65). We did brain extraction using SynthStrip (66) in Freesurfer (http://surfer.nmr.mgh.harvard.edu/) to ensure optimal co-registration. We created a T1-weighted group template and normalised to the Montreal Neurological Institute (MNI) space (1 mm³ voxel; MNI 152 template) using Advanced Normalization Tools (ANTs, http://stnava.github.io/ANTs/).

We upsampled T1 structural images in participant brain space (after removing the background noise) by a factor 2 ([0.375 × 0.375 × 0.375] mm^3^). Our purpose was to avoid losing in-plane resolution when registering the LC slab to the T1 image. This up-sampling was done using the nii_scale_dims function from an extension of SPM12 (https://github.com/rordenlab/spmScripts). We registered each participant’s MT-TFL image to the T1 image we upsampled before. This process was done in two steps: (a) To make an approximate transformation first, we manually registered the images and extracted its parameters using ITK-SNAP (67). (b) We used these parameters and performed automatic affine registration with ANTs. This allowed a precise registration at the end. Two expert raters delineated the LC looking at hyperintense values on registered MT-TFL images. The intersection between the two delineated LCs would be the final LC mask for each individual.

Echo-planar imaging (EPI) volumes were auto-reoriented by aligning the first image of the time series to a standard orientation. Static field inhomogeneities were corrected by calculating voxel displacement maps (VDMs) using the SPM FieldMap toolbox, where phase and magnitude images acquired as B0 field maps were processed to generate VDMs with parameters tailored to 7T acquisition using a customized defaults file. The EPI images were then realigned to the mean functional image and unwarped using the computed VDMs to account for both head motion and susceptibility-induced distortions, implemented via the Realign & Unwarp module in SPM12. Skull stripping was performed using SynthStrip, a deep-learning-based tool, on the mean EPI image, and the resulting brain mask was applied to the entire functional time series using FSL’s *fslmaths*. Finally, brain-extracted images were smoothed with a 3 mm full-width at half-maximum (FWHM) isotropic Gaussian kernel using SPM’s smoothing function.

### Quantification and Statistical Analysis

First-level statistical analyses were conducted within the general linear model (GLM) framework implemented in SPM12. The onsets of emotional and neutral prosody stimuli were modeled as stick functions and convolved with the canonical hemodynamic response function (HRF). Low-frequency drifts were removed using a 256 s high-pass filter. Physiological (cardiac and respiratory), estimated using the PhysIO Toolbox (Translational Neuromodeling Unit, ETH Zurich, Switzerland), were included as covariates of no interest to account for physiological noise (68).

For each participant, separate regressors were used to model the emotional task blocks or events under each of the five light conditions (0, 0.16, 37, 92, and 190 melanopic EDI). The resulting beta estimates reflected the condition-specific BOLD responses. Individual contrast images for each condition were carried forward to second-level (group) analyses. At the group level, contrast images from each participant were entered into second-level analyses using one-sample (zero-mean) *t*-tests in SPM12 to assess whether average contrast values differed significantly from zero across participants. Whole-brain voxel-wise significance was assessed at *p* < 0.05 (uncorrected). Group-level whole-brain analyses (second-level) were conducted using SPM12 and were used for visualization purposes only. All reported statistical significance levels and effect sizes are derived from the GLMMs performed on the beta coefficients extracted from the independent ROI masks.

In the subsequent post hoc analysis, we estimated the responses to the stimuli under each light condition. Separate regressors modelled each task’s block or event type under each light condition (0, 0.16, 37, 92, 190 melanopic EDI). The contrasts of interest consisted of the main effects of each regressor.

Regression beta coefficients were extracted using the REX Toolbox based on LC and 4^th^ ventricle segmentation masks, as well as anatomical atlases for the hypothalamus (https://web.mit.edu/swg/software.htm; (69)). The hypothalamus was segmented into five subregions—inferior anterior, superior anterior, inferior tubular, superior tubular, and posterior—using an automatic computational method within the 1 mm³ MNI152 template space (70). The amygdala was segmented into ten subregions based on a high-resolution probabilistic atlas (71). These regions included the lateral, basal, accessory basal, central, cortical, medial, and paralaminar nuclei, as well as the anterior amygdaloid area and the cortico-amygdaloid transition area. Statistical analyses of the activity of each region of interest (LC, hypothalamus, and amygdala subparts) were computed in R (version 4.0.8) using the afex package (Analysis of Factorial Experiments), implemented via a custom laboratory interface (FlexibleGLMM; https://puneet-talwar.shinyapps.io/FlexibleGLMM/). The analyses consisted of (2-sided) Generalized Linear Mixed Models (GLMM) with the subject included as a random factor (intercept) to account for the repeated-measures design.

The first main analysis assessed the average activity estimates to emotional and neutral stimuli independent of light in the LC. Afterwards, main analysis focusing on light exposure impact used a linear parametric modulation of the changes in activity with increasing illuminance to grasp the overall impact of illuminance change. Since the latter may have missed non-linear changes in activity with changes in illuminance, a factorial GLMM was computed, including Light (5 levels), Task (2 levels), and their interaction as fixed effects. Prior to this second phase of analysis, outlier detection was performed using Mahalanobis distance with a conservative p < .01 threshold (0.99 confidence interval); this resulted in the removal of 9 data points from the initial sample of 350. Models where covariated with age, gender, BMI and a seasonal covariate covariate was included consisting of the cosine value of the day of the year expressed in degrees of the 360-day year (December 21st = 0°; Mean = −0.094, SD = 0.63).

A final set of exploratory GLMMs was conducted to bridge regional brain activity with physiology and behavior. We first assessed the direct impact of illuminance on behavioral and physiological metrics. Two separate models were computed using Emotional Task Performance (Accuracy/RT) and Transient Pupil Dilation as dependent variables. These models included Light, Task, and their interaction as fixed effects, while controlling for Age, Gender, BMI, and Season (**Supplementary Table S3**). To investigate the functional role of LC, we conducted two additional sets of analyses. First, we tested whether LC Blood Oxygen Level-Dependent (BOLD) activity predicted behavioral performance or pupil dilation in separate models (**Supplementary Table S6**). Second, we explored potential relationship between subcortical regions and the LC. We used GLMs to determine whether activity in specific Hypothalamus (subregions 1 and 3) and Amygdala (subregions 3 and 5) clusters predicted LC BOLD activity across both neutral and emotional tasks (**Supplementary Table S7**). All models in this set were adjusted for Age, Gender, BMI, and Season to ensure the specificity of the associations.

Optimal sensitivity and power analyses in GLMMs remain under investigation (72). We nevertheless computed a prior sensitivity analysis to determine the minimum detectable effect size in our main analyses, given our sample size. According to G*Power 3 (version 3.1.9.7, (73)), a sample of N = 35 (with power = 0.80 and α = 0.05) allowed us to detect effect sizes of f^2^ > 0.31 (R^2^ > 0.23; r > 0.48) within a linear multiple regression framework including two tested predictors (illuminance and stimulus type) and up to four covariates (age, sex, BMI, and season).

## Acknowledgments

The study was conducted at the GIGA-In Vivo Imaging technological platform of ULiège, Belgium. The authors thank Christine Bastin, Nikita Beliy, Annick Claes, Christian Degueldre, Catherine Hagelstein, Gregory Hammad, Brigitte Herbillon, Ekaterine Koshmanova, Benjamin Lauricella, and Pierre Maquet for their help over the different steps of the study. In particular We also thank Anthony Harry for the preparation of Figure 1.

It was supported by the Belgian Fonds National de la Recherche Scientifique (FNRS; CDR J.0222.20, & J.0216.24), the European Union’s Horizon 2020 research and innovation program under the Marie Skłodowska-Curie grant agreement No 860613, the Fondation Léon Frédéricq, ULiège - U. Maastricht Imaging Valley, Fondation Recherche Alzheimer (SAO-FRA 2019/0025 & 2022/0014), the European Regional Development Fund (Biomed-Hub, WALBIOIMAGING), and Siemens. None of these funding sources had any impact on the design of the study nor the interpretation of the findings.

EB was supported by the Maastricht University - Liège University Imaging Valley. RS and FB were supported by the European Union’s Horizon 2020 research and innovation program under the Marie Skłodowska-Curie grant agreement No 860613. IC, IP, NM, and GV are/were supported by the FRS-FNRS. PT and LL are/were supported by the EU Joint Programme Neurodegenerative Disease Research (JPND) IRONSLEEP and SCAIFIELD projects, respectively – FNRS references: PINT-MULTI R.8011.21 & R.8006.20). LL is supported by the European Regional Development Fund (WALBIOIMAGING). MZ is supported the Fondation Recherche Alzheimer (SAO-FRA 2022/0014). HILJ is funded through the National Institutes of Health (R01AG062559, R01AG068062, R01AG082006, was chair and is currently the immediate past chair of the Neuromodulatory Subcortical Systems Professional Interest Area of ISTAART and served as an advisory board member of ISTAART.

