## Supplementary Material for "*In vivo* modulation of locus coeruleus activity by light"

**Supplementary Table S1. Demographics of study sample.**

|  | **Total Sample** |
| --- | --- |
| **Number of Participants** | 35 |
| **Age** | 24.49 ± 3.32 |
| **Sex (M)** | 13 |
| **Mood (BDI-II)** | 6.93 ± 5.84 |
| **Anxiety (BAI)** | 5.37 ± 4.22 |
| **Sleep quality (PSQI)** | 4.17 ± 2.55 |
| **Seasonality (SPAQ)** | 1.1 ± 0.8 |
| **Chronotype (HO)** | 46.52 ± 8.93 |
| **Daytime sleepiness (ESS)** | 6.58 ± 3.07 |
| **Years of Education** | 14.7 ± 3.15 |
| **Sleep duration (night before fMRI protocol – sleep diary based)** | 7.51 ± 0.85 |
| **Number of participants per month** | j:1; f:2; m:2; a:5; m:4; j:3; j:1; a:3; s:5; o:4; n:4; d:1 |
| **(January to December)** |  |

Total number of participants included in the analyses. BDI-II, Beck's Depression Inventory; BAI, Beck Anxiety Inventory; PSQI, Pittsburgh Sleep Quality Index; SPAQ, Seasonal Pattern Assessment Questionnaire; HO, Horne and Östberg; ESS, Epworth Sleepiness Scale. Refer to the method for the references to the scales and questionnaires.

**Supplementary Table S2. Light characteristics.**

|  | **Low BEL** | **Mid BEL** | **High BEL** | **Orange** |
| --- | --- | --- | --- | --- |
| **Lux** | 47 | 116 | 240 | 7.5 |
| **Peak Spectral Irradiance (nm)** | 460 | 460 | 460 | 590 |
| **Melanopic EDI (lux; ipRGCs)** | 37 | 92 | 190 | 0.16 |
| **Rhodopic EDI (lux; Rods)** | 39 | 97 | 201 | 0.94 |
| **Cyanopic EDI (lux; S-cones)** | 32 | 79 | 163 | 0 |
| **Chloropic EDI (lux; M-cones)** | 44 | 110 | 227 | 5 |
| **Erythropic EDI (lux ; L-cones)** | 46 | 113 | 233 | 8 |
| **Irradiance (µW/cm²)** | 15 | 36 | 75 | 1.4 |
| **Photon flux(1/cm²/s)** | 4.12E+13 | 1.02E+14 | 2.10E+14 | 4.24E+12 |
| **Log Photon Flux (log₁₀ (1/cm²/s)** | 13.61 | 14.01 | 14.32 | 12.63 |
| **Narrowband peak** | - | - | - | 589 |
| **Narrowband FWHM** | - | - | - | 10 |

Detailed characteristics of the four conditions used in fMRI protocol. Blue enriched (BEL) (low, mid, and high) and monochromatic (589nm). ipRGCs: intrinsically photosensitive retinal ganglion cells. FWHM: full width at half maximum.

**Supplementary Table 3. Effect of illuminance on performance to the emotional task and transient pupil dilation.** F-statistics are reported for fixed effects with degrees of freedom (df) in parentheses, estimated using the Satterthwaite approximation. *P < .05, **P < .01, ***P < .001.

| **Emotional Task Performance with Light** | | | |
| --- | --- | --- | --- |
| **Effect** | ***F* (*df*)** | ***p*** | ***η2*** |
| **Fixed Effects** | | | |
| **Light** | 2.41 (4,92) | 0.054 | 0.095 |
| **Task** | 1.97 (1,19) | **<0.001***** | 0.405 |
| **Age** | 0.53 (1,19) | 0.176 | 0.094 |
| **Sex** | 1.37 (1,19) | 0.475 | 0.027 |
| **BMI** | 0.11 (1,19) | 0.256 | 0.067 |
| **Season** | 15.66 (1,23) | 0.747 | 0.005 |
| **Light x Task** | 0.39 (4,92) | 0.816 | 0.017 |
| **Emotional Transient Pupil Dialation with Light** | | | |
| **Effect** | ***F* (*df*)** | ***p*** | ***η2*** |
| **Fixed Effects** | | | |
| **Light** | 9.2347 (4, 90.64) | **0.003 ***** | 0.290 |
| **Age** | 0.1834 (1, 18.98) | 0.673 | 0.010 |
| **Sex** | 3.0432 (1, 18.87) | 0.097 | 0.139 |
| **BMI** | 0.1170 (1, 18.73) | 0.736 | 0.006 |
| **Season** | 0.1460 (1, 18.73) | 0.707 | 0.008 |
| **Task** | 1.5519 (1, 22.83) | 0.225 | 0.064 |
| **Light x Task** | 0.2904 (4, 90.52) | 0.884 | 0.013 |

**Supplementary Table S4. Post hoc contrasts of the GLMM considering the activity of the LC under each illuminance (related to table 2 of the main text)**

| **Condition** | **Comparison** | ***t*** | ***p (uncorrected)*** |
| --- | --- | --- | --- |
| **0 lux** | Neut - Emo | 0.125 | 0.9006 |
| **0.16 lux** | Neut - Emo | 0.206 | 0.8372 |
| **37 lux** | Neut - Emo | -0.443 | 0.6583 |
| **92 lux** | Neut - Emo | -1.342 | 0.1815 |
| **190 lux** | Neut - Emo | -4.512 | **<0.001***** |
| **Neut** | Darkness (0 lux) - Orange (0.16 lux) | -0.474 | 0.6357 |
| **Neut** | Darkness (0 lux) - B1 (37 lux) | -0.22 | 0.8926 |
| **Neut** | Darkness (0 lux) - B2 (92 lux) | -0.260 | 0.7953 |
| **Neut** | Darkness (0 lux) - B3 (190 lux) | -1.548 | 0.1229 |
| **Neut** | Orange (0.16 lux) - B1 (37 lux) | 0.605 | 0.5457 |
| **Neut** | Orange (0.16 lux) - B2 (92 lux) | 0.219 | 0.8266 |
| **Neut** | Orange (0.16 lux) - B3 (190 lux) | -1.046 | 0.2965 |
| **Neut** | B1 (37 lux) - B2 (92 lux) | -0.393 | 0.6950 |
| **Neut** | B1 (37 lux) - B3 (190 lux) | -1.670 | 0.0963 |
| **Neut** | B2 (92 lux) - B3 (190 lux) | -1.288 | 0.1989 |
| **Emo** | Darkness (0 lux) - Orange (0.16 lux) | -0.545 | 0.5468 |
| **Emo** | Darkness (0 lux) - B1 (37 lux) | 0.621 | 0.5350 |
| **Emo** | Darkness (0 lux) - B2 (92 lux) | 0.997 | 0.3200 |
| **Emo** | Darkness (0 lux) - B3 (190 lux) | 2.431 | **0.0158*** |
| **Emo** | Orange (0.16 lux) - B1 (37 lux) | 1.149 | 0.2516 |
| **Emo** | Orange (0.16 lux) - B2 (92 lux) | 1.516 | 0.1309 |
| **Emo** | Orange (0.16 lux) - B3 (190 lux) | 2.916 | **0.004**** |
| **Emo** | B1 (37 lux) - B2 (92 lux) | 0.372 | 0.7101 |
| **Emo** | B1 (37 lux) - B3 (190 lux) | 1.793 | 0.0742 |
| **Emo** | B2 (92 lux) - B3 (190 lux) | 1.424 | 0.1559 |

**Supplementary Table S5. 4^th^ Ventricle Main Stats.** F-statistics are reported for fixed effects with degrees of freedom (df) in parentheses, estimated using the Satterthwaite approximation. *P < .05, **P < .01, ***P < .001.

| **Lightmod Model** | | | |
| --- | --- | --- | --- |
| **Emotional Task 4th Ventricle Analysis** | | | |
| **Effect** | ***F* (*df*)** | ***p*** | ***η2*** |
| **Task** | 1.90 (1,34) | 0.177 | 0.053 |
| **Age** | 1.83 (1,30) | 0.186 | 0.058 |
| **Sex** | 0.34 (1,30) | 0.564 | 0.011 |
| **BMI** | 0.07 (1,30) | 0.793 | 0.002 |
| **Season** | 0.46 (1,30) | 0.502 | 0.015 |

**Supplementary Table 6. Association between performance to the task and pupil transient dilation to the emotional task and the activity of LC for each illuminance.** F-statistics are reported for fixed effects with degrees of freedom (df) in parentheses, estimated using the Satterthwaite approximation. *P < .05, **P < .01, ***P < .001.

| **Effect** | ***F* (*df*)** | ***p*** | ***η2*** |
| --- | --- | --- | --- |
| **Emotional Task Performance with LC** | | | |
| **Light** | 2.292 (4, 211.10) | .061 | 0.042 |
| **Age** | 1.982 (1, 18.99) | .175 | 0.095 |
| **Sex** | 0.518 (1, 19.01) | .480 | 0.027 |
| **BMI** | 1.373 (1, 18.98) | .256 | 0.067 |
| **Season** | 0.110 (1, 18.98) | .744 | 0.006 |
| **LC** | 0.044 (1, 217.63) | .834 | 0.000 |
| **Effect** | ***F* (*df*)** | ***p*** | ***η2*** |
| **Emotional Transient Pupil Dialation with LC** | | | |
| **Light** | 10.219 (4, 206.42) | **<.001***** | 0.165 |
| **Age** | 0.198 (1, 19.03) | .661 | 0.010 |
| **Sex** | 3.001 (1, 18.87) | .100 | 0.137 |
| **BMI** | 0.110 (1, 18.96) | .744 | 0.006 |
| **Season** | 0.150 (1, 18.79) | .703 | 0.008 |
| **LC** | 0.006 (1, 212.73) | .938 | 0.000 |

**Supplementary Table 7. Association between LC activity and amygdala/hypothalamus subparts (related to Figure 3)**

| **Task** | **Predictor Region** | ***t*** | ***p*** | ***η2*** |
| --- | --- | --- | --- | --- |
| **LC activity and Hypothalamus and Amygdala Relationships** | | | | |
| **Neutral** | Anterior Inferior Hypothalamus | 1.756 | 0.09 | 0.154 |
|  | Posterior Lateral Hypothalamus | -0.388 | 0.701 | 0.069 |
|  | Basomedial Nuclei Amygdala | -0.749 | 0.460 | 0.082 |
|  | Medial and Cortical Nuclei Amygdala | 0.807 | 0.426 | 0.085 |
| **Emotional** | Anterior Inferior Hypothalamus | 0.059 | 0.953 | 0.072 |
|  | Posterior Lateral Hypothalamus | 2.181 | **0.037 *** | 0.203 |
|  | Basomedial Nuclei Amygdala | 2.080 | **0.046 *** | 0.192 |
|  | Medial and Cortical Nuclei Amygdala | 1.431 | 0.163 | 0.133 |

General Linear Models (GLM) examining subcortical predictors of Locus Coeruleus (LC) BOLD activity across Neutral and Emotional tasks. Relationships were examined using General Linear Models (GLM). To ensure the specificity of these associations, Age, Gender, BMI, and Season were included as covariates in all models to account for potential inter-individual variability and seasonal influences on LC and subcortical activity. Steiger’s Z-tests revealed no significant differences in LC-association strength between subparts within tasks (p > 0.19). *P < .05, **P < .01, ***P < .001.

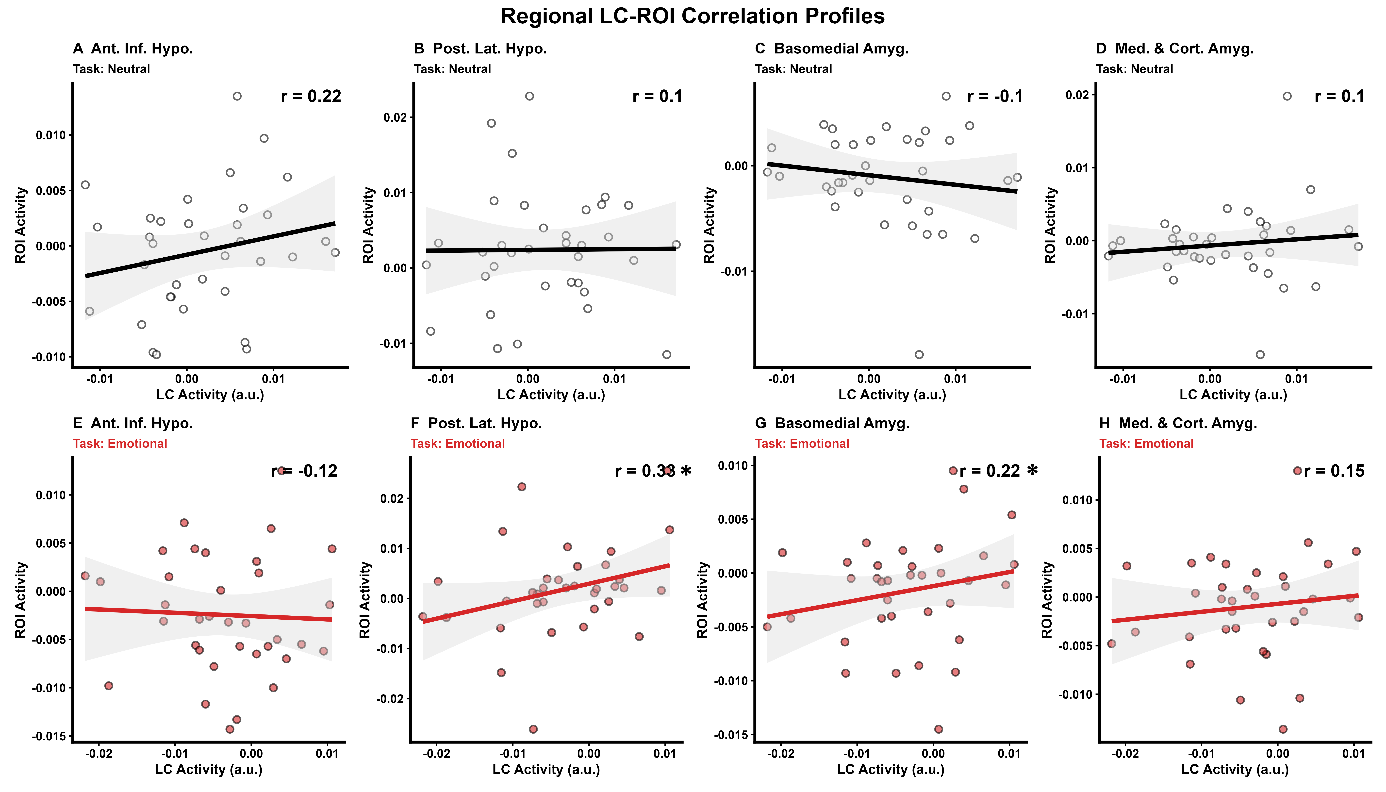

**Supplementary Figure 1. Regional LC-ROI Correlation Profiles.** This eight-panel scatterplot matrix displays the relationship between Locus Coeruleus (LC) activity (a.u.; horizontal axes) and the activity of four specific brain Regions of Interest (ROIs; vertical axes): Anterior Inferior Hypothalamus (Ant. Inf. Hypo.), Posterior Lateral Hypothalamus (Post. Lat. Hypo.), Basomedial Amygdala (Basomedial Amyg.), and Medial & Cortical Amygdala (Med. & Cort. Amyg.). Panels **A–D** (top row, blue headers) illustrate correlations during the Neutral task condition, using black regression lines and open circles. Panels **E–H** (bottom row, red headers) show correlations during the Emotional task condition, utilizing red regression lines and red-filled circles. Spearman correlation coefficients (*r*) are displayed in the upper right of each panel; l; asterisks indicate significant correlations at p < .05. Bold, consistent lines signify mean trends, and the light grey shaded ribbons indicate the 95% confidence interval for each correlation. Data were processed to map discrete variables onto continuous plotting spaces and exclude outliers beyond strict axis limits for consistent visualization. (refer to Suplementary Table 7 for full statistics).

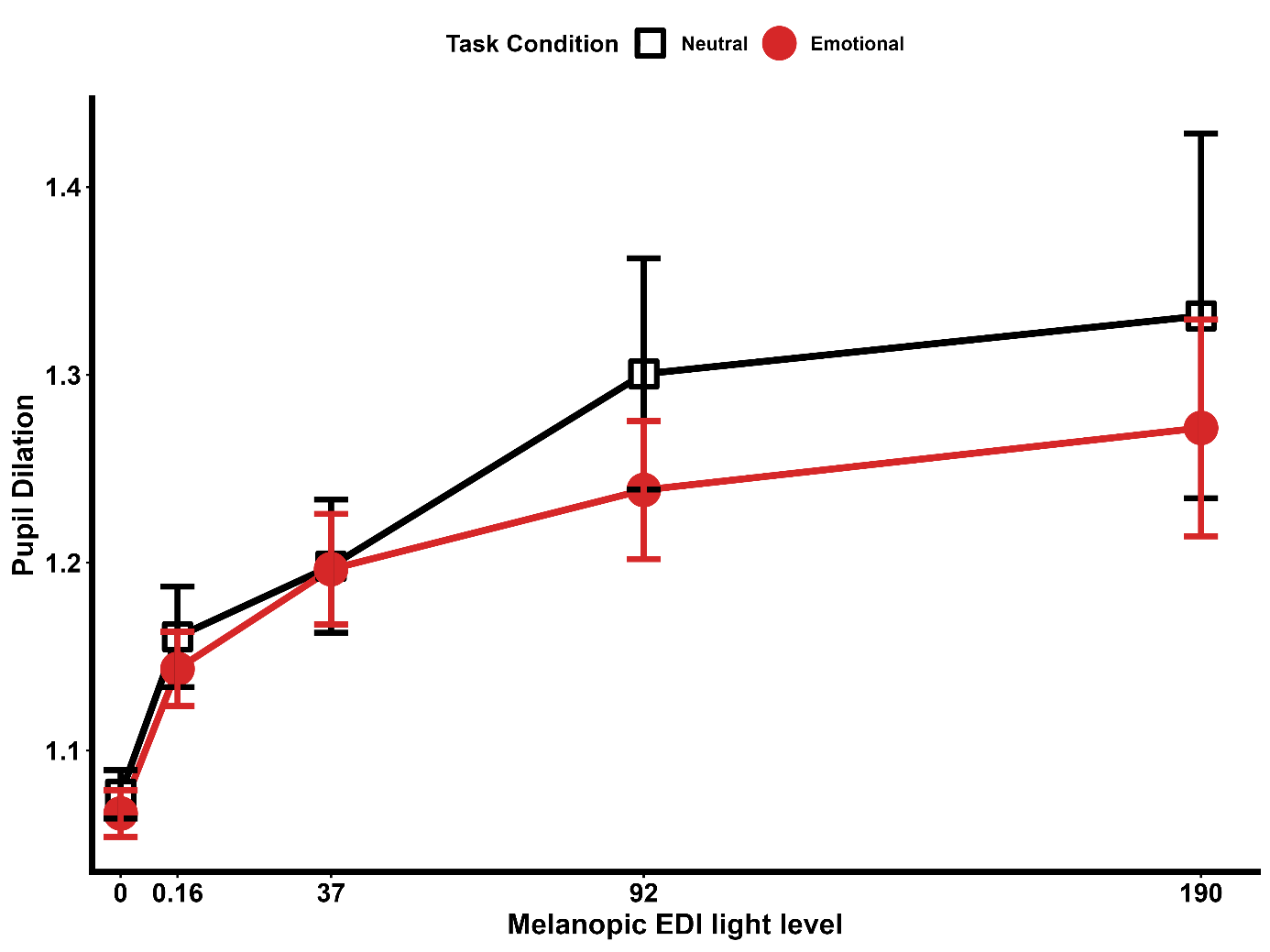

**Supplementary Figure 2. Effect of Light Intensity and Task Condition on Pupil Dilation.** This line graph illustrates the mean pupil dilation (ratio, a.u.) across five Melanopic EDI light levels (0, 0.16, 37, 92, and 190). Data are stratified by task condition: **Neutral** (black line, open squares) and **Emotional** (red line, filled circles). Error bars represent the standard error of the mean (SEM). The x-axis is plotted on a non-linear scale to reflect specific experimental light intensity intervals. Statistical analysis revealed a significant main effect of Light (p = .003), indicating a dose-dependent increase in pupil dilation as light intensity increases (refer to Supplementary Table 3 for full statistics).
